# TMEM16A activation for the fast block to polyspermy in the African clawed frog does not require conventional activation of egg PLCs

**DOI:** 10.1101/2022.08.30.505853

**Authors:** Kayla M. Komondor, Rachel E. Bainbridge, Katherine G. Sharp, Joel C. Rosenbaum, Anne E. Carlson

**Author notes:** These authors contributed equally to this work (Kayla M. Komondor, Rachel E. Bainbridge). Correspondence to: Anne E. Carlson.

## Abstract

Fertilization of an egg by more than one sperm, a condition known as polyspermy, leads to gross chromosomal abnormalities and is embryonic lethal for most animals. Consequently, eggs have evolved multiple processes to stop supernumerary sperm from entering the nascent zygote. For external fertilizers, such as frogs and sea urchins, fertilization signals a depolarization of the egg membrane, which serves as the fast block to polyspermy. Sperm can bind to, but will not enter, depolarized eggs. In eggs from the African clawed frog, *Xenopus laevis*, the fast block depolarization is mediated by the Ca^2+^ activated Cl^−^ channel TMEM16A. To do so, fertilization activates a phospholipase C, which generates IP_3_ to signal a Ca^2+^ release from the ER. Currently, the signaling pathway by which fertilization activates PLC remains unknown. Here, we sought to uncover this pathway by targeting the canonical activation of the PLC isoforms present in the *X. laevis* egg: PLCγ and PLCβ. We observed no changes to the fast block in *X. laevis* eggs inseminated in inhibitors of tyrosine phosphorylation, used to stop activation of PLCγ, or inhibitors of G_αq/11_ pathways, used to stop activation of PLC_β_. These data suggest that the PLC that signals the fast block depolarization in *X. laevis* is activated by a novel mechanism.

## Introduction

For most sexually reproducing animals, only eggs fertilized by one sperm can successfully initiate embryonic development (Hassold et al., 1980; Rojas et al., 2021). Fertilization of an egg by more than one sperm, a condition known as polyspermy, is lethal to the developing embryo (Wong and Wessel, 2006). To prevent these catastrophic consequences, eggs have evolved various processes called polyspermy blocks that stop sperm from entering already-fertilized eggs (Wong and Wessel, 2006; Evans, 2020; Fahrenkamp et al., 2020). The molecular details of these processes are still being uncovered.

Two common polyspermy blocks are named based on their relative timing: the “fast” and “slow” blocks to polyspermy (Jaffe and Gould, 1985; Bianchi and Wright, 2016). The eggs of nearly all sexually reproducing animals use the slow block to polyspermy, which occurs minutes after fertilization and involves the exocytosis of cortical granules from the egg and establishment of a physical barrier around the nascent zygote (Wong and Wessel, 2006; Evans, 2020). Eggs from externally fertilizing animals also use the fast block to polyspermy (Wozniak and Carlson, 2020). During the fast block to polyspermy, fertilization immediately activates a depolarization of the egg plasma membrane (Jaffe, 1976). This membrane potential change allows sperm to bind to, but not penetrate the egg (Jaffe, 1976). We are still uncovering the molecular pathways that underlie this process (Tembo et al., 2020; Wozniak and Carlson, 2020). Here we sought to uncover how fertilization signals the fast block in eggs from the African clawed frog, *Xenopus laevis*.

We have previously demonstrated that fertilization of *X. laevis* eggs opens the Ca^2+^-activated Cl^−^ channel TMEM16A (Wozniak et al., 2018a), allowing Cl^−^ to leave the cell and depolarize its membrane. TMEM16A channels are activated by elevated cytoplasmic Ca^2+^, and fertilization opens these channels via phospholipase C (PLC) mediated activation of the inositol trisphosphate (IP_3_) receptor on the endoplasmic reticulum (ER), to enable Ca^2+^ release from ER (Wozniak et al., 2018b). Notably, both the fast and slow polyspermy blocks in *X. laevis* eggs require PLC activation and IP3-signaled Ca^2+^ release from the ER (Nuccitelli et al., 1993; Fontanilla and Nuccitelli, 1998; Wozniak et al., 2018b).

Conflicting data has been published regarding how fertilization in *X. laevis* activates egg PLCs. For example, by imaging cytoplasmic Ca^2+^, Glahn and colleagues reported that injecting *X. laevis* eggs with tyrosine kinase inhibitors lavendustin A or tyrphostin B46 stopped the fertilization evoked Ca^2+^ wave (Glahn et al., 1999). Similarly, Sato and colleagues reported that inhibiting tyrosine phosphorylation with 100 μM genistein completely prevented embryonic development of *X. laevis* eggs (Sato et al 1998). Further work suggested that fertilization in *X. laevis* eggs activates PLCγ via tyrosine phosphorylation and that PLCγ specifically was necessary for the Ca^2+^ increase in the egg following fertilization (Sato et al., 2000). These experiments gave rise to the hypothesis that sperm bind to the extracellular domain of uroplakin III, which signals activation of a Src family kinase which then tyrosine phosphorylates PLCγ (Mahbub Hasan et al., 2005). However, Runft and colleagues imaged cytoplasmic Ca^2+^ in *X. laevis* eggs and reported that stopping either PLCγ activity by overexpressing the Src-homology 2 (SH2) domain or PLCβ activation with a Gaq targeting antibody, failed to prevent the fertilization-signaled increase in cytoplasmic Ca^2+^ (Runft et al., 1999). These data suggest that neither tyrosine phosphorylation of PLCγ nor Gα_q_ activation of PLCβ are needed for the fast block.

Here, we sought to uncover the connection between fertilization and PLC activation during the fast block in *X. laevis* eggs. From analysis of existing transcriptomics and proteomics datasets, we identified three PLC isoforms in *X. laevis* eggs: PLCγ1, PLCβ1 and PLCβ3. We then determined which of the signaling pathways upstream of each PLC subtype were required for the fast block in *X. laevis*. We report that the canonical activation pathways for neither PLCγ1 nor PLCβare required for the fast block in *X. laevis* eggs. Our findings indicate that PLC activation during *X. laevis* fertilization proceeds through a new, yet to be determined pathway.

## Materials and methods

### Reagents

Genistein was obtained from Alfa Aesar (Thermo Fisher Scientific; Tewksbury, MA), Lavendustin A/B were obtained from Santa Cruz Biotechnology (Dallas, TX), and human chorionic gonadotropin was purchased from Covetrus (Dublin, OH). Leibovitz’s-15 (L-15) medium (without L-glutamine) was purchased from Sigma-Aldrich. All other materials, unless noted, were purchased from Thermo Fisher Scientific.

### Solutions

Modified Ringer’s (MR) solution was used as the base for all fertilization and recording assays in this study (100 mM NaCl, 1.8 mM KCl, 2.0 mM CaCl_2_, 1.0 mM MgCl_2_, 5.0 mM HEPES, pH 7.8). The MR solution was filtered using a sterile, 0.2 μm polystyrene filter (Heasman et al., 1991) and diluted for experimentation as follows: Fertilization recordings were performed in 20% MR (MR/5) with or without indicated inhibitors. Following whole cell recordings, developing embryos were stored and monitored in 33% MR (MR/3).

OR2 and ND96 solutions were used for collection and storage of the immature oocytes. OR2 (82.5 mM NaCl, 2.5 mM KCl, 1 mM MgCl_2_, 5 mM HEPES, pH 7.6) and ND96 (96 mM NaCl, 2 mM KCl, 1.8 mM CaCl_2_, 1 mM MgCl_2_, 5 mM HEPES, 5 mM sodium pyruvate, gentamycin, pH 7.6) were filtered using a sterile, 0.2 μM polystyrene filter.

MR solutions containing inhibitors were prepared from concentrated stock solutions made in DMSO. Final DMSO content was maintained below 2% solution, a concentration that does not interfere with the fast block (Wozniak et al., 2018b).

### Animals

All animal procedures were conducted using acceptable standards of humane animal care and approved by the Animal Care and Use Committee at the University of Pittsburgh. *X. laevis* adults were obtained commercially from Nasco or Xenopus 1 and housed at 20°C with a 12-h/12-h light/dark cycle.

### Collection of gametes

#### Eggs

To obtain fertilization-competent eggs, sexually mature *X. laevis* females were induced to ovulate via injection of 1,000 IU human chorionic gonadotropin into the dorsal lymph sac and overnight incubation at 14-16°C for 12-16 hours (Wozniak et al., 2017). Females typically begin to lay eggs within 2 hours after being moved to room temperature. Eggs were collected on dry Petri dishes and used within 10 minutes of laying.

#### Sperm

To obtain sperm, testes were harvested from sexually mature *X. laevis* males (Wozniak et al., 2017). Males were euthanized by a 30-minute immersion in 3.6 g/L tricane-S (MS-222), pH 7.4, before testes were harvested and cleaned. Testes were then stored at 4°C in MR for use on the day of the dissection, or in L15 Leibovitz’s Medium without glutamine for use up to 1 week later.

#### Oocytes

*X. laevis* oocytes were collected by procedures described in previous manuscripts (Tembo et al., 2019). Briefly, ovarian sacs were obtained from *X. laevis* females anesthetized with a 30-minute immersion in 1.0 g/L tricaine-S (MS-222) at pH 7.4. Ovarian sacs were manually pulled apart and incubated for 90 minutes in 1 mg/mL collagenase in ND96 supplemented with 5 mM sodium pyruvate and 10 mg/L of gentamycin. Collagenase was removed by repeated washes with OR2, and healthy oocytes were sorted prior to storage at 14 °C in ND96 supplemented with sodium pyruvate and gentamycin.

### Sperm preparation and *in vitro* fertilization

For *in vitro* fertilizations during whole cell recordings, sperm suspensions were made by macerating 1/10 of the *X. laevis* testis in 200 μL MR/5. This solution was kept at 4°C for up to 1 hour for use. Up to three sperm additions were added to each egg while recording, with approximately 10 minutes between additions. To monitor for development after recording, eggs inseminated during whole cell recordings were transferred to MR/3 and incubated at room temperature for 2 hours. Development was assessed based on the appearance of cleavage furrows (Wozniak et al., 2017).

### Electrophysiology

Electrophysiological recordings were made using TEV-200A amplifiers (Dagan Co.) and digitized by Axon Digidata 1550A (Molecular Devices). Data were acquired with pClamp Software (Molecular Devices) at a rate of 5 kHz. Pipettes used to impale the *X. laevis* eggs for recordings were pulled from borosilicate glass for a resistance of 5-15 MΩ and filled with 1 M KCl.

#### Whole cell recordings

Resting and fertilization potentials were quantified ~10 s before and after the depolarization, respectively. Depolarization rates of each recording were quantified by determining the maximum velocity of the quickest 1-mV shift in the membrane potential (Wozniak et al., 2018a).

#### Two electrode voltage clamp recordings

The efficacy of inhibitors targeting PLCβ activation was screened by recording xTMEM16A currents in the two-electrode voltage clamp configuration on *X. laevis* oocytes clamped at −80 mV. Blue/green light was applied using a 250 ms exposure to light directed from the opE-300^white^ LED Illumination System (CoolLED Ltd) and guided by a liquid light source to the top of the oocytes in 35 mm petri dishes. Background-subtracted peak currents were quantified from two consecutive recordings (one before and one during application of screened inhibitors). The proportional difference between peak currents before and with inhibitor for each oocyte was used to quantify inhibition.

### Polyspermy assay

To determine the incidence of polyspermy with or without inhibitors, inseminated eggs were kept for observation for 2 hours following whole cell recordings. Successful monospermic fertilization was defined by symmetrical patterns of cleavage furrows in embryos, while polyspermic fertilization was defined by asymmetric patterns of cleavage furrows (Elinson, 1975; Grey et al., 1982).

### Exogenous protein expression in *X. laevis* oocytes

The cDNAs encoding the platelet derived growth factor receptor (PDGF-R) (Gagoski et al., 2016) or the rhodopsin-muscarinic receptor type 1 chimera (opto-M1R) (Morri et al., 2018) were purchased from Addgene (plasmids 67130 and 106069 respectively) and were engineered into the GEMHE vector using overlapping extension PCR. The sequences for all constructs were verified by automated sequencing (Gene Wiz or Plasmidsaurus). The cRNAs were transcribed using the T7 mMessage mMachine (Ambion). Defolliculated oocytes were injected with cRNA and used 3 days following injection.

### Removal of egg jelly

For procedures requiring the removal of jelly, eggs were incubated at room temperature for 5 minutes before insemination with sperm suspension prepared as described previously (Wozniak et al., 2020). Activated eggs, identified by their ability to roll so the animal pole faced up and displayed a contracted animal pole, were used for western blot preparations. To remove the jelly, eggs were placed on agar in a 35 mm petri dish, in MR/3 with 45 mM β-mercaptoethanol (BME), pH 8.5. During the BME incubation, eggs were gently agitated for 1-2 minutes, until their external jelly visibly dissolved, as indicated by close nestling of the eggs and loss of visible jelly. To remove the BME solution, de-jellied eggs were then moved with a plastic transfer pipette to a petri dish coated with 1% agar in MR/3 pH 6.5 and agitated for an additional minute. Eggs were then transferred three times to additional agar coated dishes with MR/3 pH 7.8, swirling gently and briefly in each.

### Sample preparation and western blot

Oocytes and eggs were lysed using a Dounce homogenizer and ice-cold oocyte homogenization buffer (OHB) (10 mM HEPES, 250 mM sucrose, 5 mM MgCl_2_, 5% glycerol supplemented with protease and phosphatase inhibitors in a 1:100 dilution) (Hill et al., 2005). 100 μL of OHB was used per 10 eggs or oocytes. Cellular debris was removed by centrifugation at 500 RCF for 5 mins at 4 °C and the resulting pellet was then resuspended in 100 μL OHB. The sample was again sedimented and the supernatant from both the successive sedimentations was pooled and again centrifuged at 18,213 RCF for 15 mins at 4 °C. The supernatant was then sedimented at 18,213 RCF for 15 mins at 4 °C. 30 μL of this supernatant was combined with 10 μL sample buffer (50 mM Tris pH 6.8, 2% SDS, 10% glycerol, 1%β-mercaptoethanol, 12.5 mM EDTA, 0.02% bromophenol blue) before incubating at 95 °C for 1 min.

Samples were resolved by electrophoresis on precast 4-12% BIS-TRIS PAGE gels (Invitrogen) run in TRIS-MOPS (50 mM MOPS, 50 mM Tris, 1 mM EDTA, 0.1% SDS) running buffer, followed by wet transfer to a nitrocellulose membrane at 10 V for 1 hour in Bolt (Invitrogen) transfer buffer. Loading was evaluated by Ponceau S. Blocking was performed for 1 hour at room temperature using Superblock (Thermo) buffer. Primary antibody incubation was performed overnight at 4 °C with anti-PLCγ[pY783] (Abcam) (1:1000) and secondary antibody was performed for 1 hour at room temperature using goat anti-rabbit HRP (Invitrogen) (1:10000). All washes were performed using TBST (20 mM Tris, 150 mM NaCl, 0.1% Tween-20, pH 8). Blots were resolved using Supersignal Pico (Pierce) on a GE/Amersham 600RGB imager using chemiluminescence settings.

### Quantification and statistical analyses

All electrophysiology recordings were analyzed with Igor (WaveMetrics) and Excel (Microsoft). Data for each experimental condition are displayed in Tukey box plot distributions, where the box contains the data between 25% and 75% and the whiskers span 10-90%. All conditions include trials that were conducted on multiple days using gametes from multiple individuals. Analysis of variance (ANOVA) with post hoc Tukey honestly significant difference test was used to determine differences between inhibitor treatments. Depolarization rates were log_10_ transformed before statistical analyses.

### Imaging

*X. laevis* eggs and embryos were imaged using a stereoscope (Leica Microsystems) equipped with a Leica 10447157 1X objective and DFC310 FX camera. Images were analyzed using LAS (version 3.6.0 build 488) software and photoshop (Adobe).

## Results

### Fertilization signals a depolarization in *X. laevis* eggs

To study the fast block to polyspermy in *X. laevis*, we performed whole cell recordings on eggs before and following sperm application. Sperm application during the recording resulted in a fertilization-signaled depolarization (Fig. 1A). For eggs inseminated under control conditions, we found that the average resting potential was −15.9 ± 1.0 mV, the potential following fertilization was 6.5 ± 2.4 mV (N=27, Fig. 1B), and the mean rate of depolarization was 4.4 ± 1.7 mV/ms (Fig. 1C). We have previously demonstrated that blocking the IP3-evoked Ca^2+^ release from the ER, with either inhibition of PLC or the IP3 receptor, was sufficient to completely stop the fast block in *X. laevis* (Fig. 1E) (Wozniak et al., 2018a). Here we similarly report that in the presence of 1 μM of the general PLC inhibitor U73122, fertilization never evoked a depolarization in *X. laevis* eggs (N=5, Fig. 1B & D). However, all 5 eggs fertilized in U73122 developed asymmetric cleavage furrows, consistent with polyspermic fertilization (Fig. 1F) (Elinson, 1975; Grey et al., 1982). These results substantiate a requirement for PLC to activate the fast block in *X. laevis* eggs.

**Figure 1.**
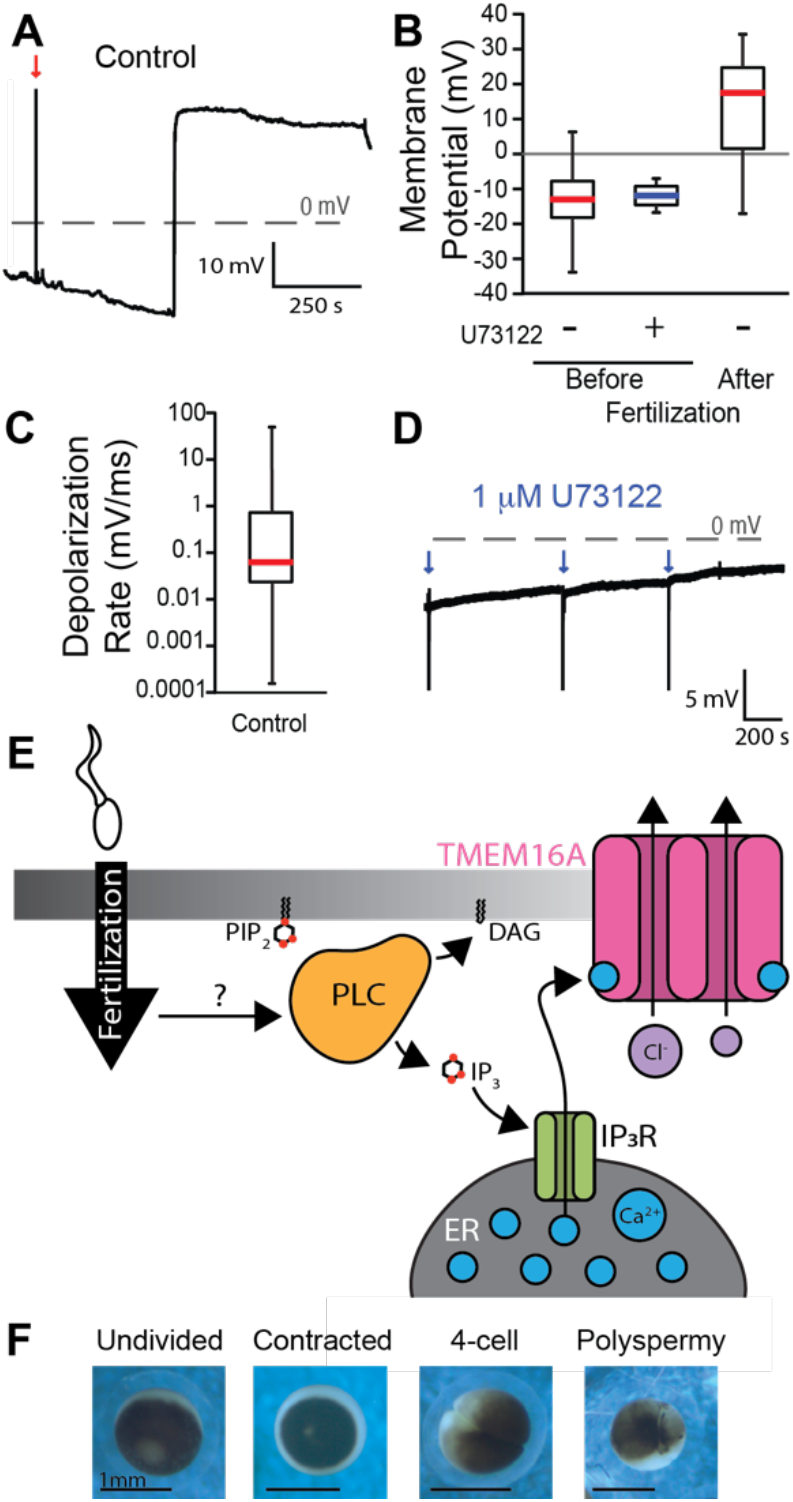
Fertilization signals a PLC-mediated depolarization in *X. laevis* eggs. **(A)** Representative whole-cell recording in control conditions (diluted modified Ringer’s solution). The red arrow indicates the time of sperm addition. Tukey box plot distributions of **(B)** resting and fertilization potentials in control conditions and resting potential in 1 mM U73122, a PLC inhibitor, and **(C)** depolarization rate in control conditions. The middle line denotes the median value, the box indicated 25-75%, and the whiskers denote 10-90%. **(D)** Representative whole-cell recording in 1 mM U73122. Blue arrows indicate sperm addition. **(E)** PIP_2_ is cleaved by phospholipase C (PLC) during the fast block to create IP_3_ and DAG. Generation of IP_3_ propagates the fast block to polyspermy. **(F)** Representative images of an undivided, 4-cell stage, and polyspermic *X. laevis* egg.

### Three PLCs in the *X. laevis* egg are candidates for triggering the fast block

We sought to uncover how fertilization activates PLC to signal the fast block of *X. laevis* eggs. Because the different PLC subtypes are activated by different signaling pathways, we first sought to identify which PLC isoforms are present in *X. laevis* eggs. To do so, we interrogated two previously published high-throughput expression datasets. First, we examined the proteome of fertilization-competent eggs (Wuhr et al., 2014) and queried for all proteins encoded by known PLC genes (Fig. 2). We found that three PLC proteins are present in the egg: PLCγ1 (encoded by the PLCG1 gene), PLCβ1 and PLCβ3 (encoded by the PLCB1 and PLCB3 genes, respectively). PLCγ1 is the most abundant in the *X. laevis* egg, present at 85.2 nM, an approximate 20-fold higher concentration than either PLCβ1 (3.9 nM), or PLCβ3 (4.3 nM). We also looked at an RNA-sequencing dataset acquired during different stages of development of *X. laevis* oocytes and eggs (Session et al., 2016). We reasoned that if these three PLC proteins are present in the fertilization-competent eggs, the RNA encoding these enzymes should be present in the developing gamete. Indeed, mRNA for all three PLC types was present in the developing oocytes (Fig. 2). We also observed RNA encoding isoforms not found in the egg (Fig. 2); this is expected as the unfertilized egg contains all of the RNA translated before the maternal-to-zygotic transition occurs in during *X. laevis* development, when the embryo is approximately 4,000 cells (Yang et al., 2015).

**Figure 2.**
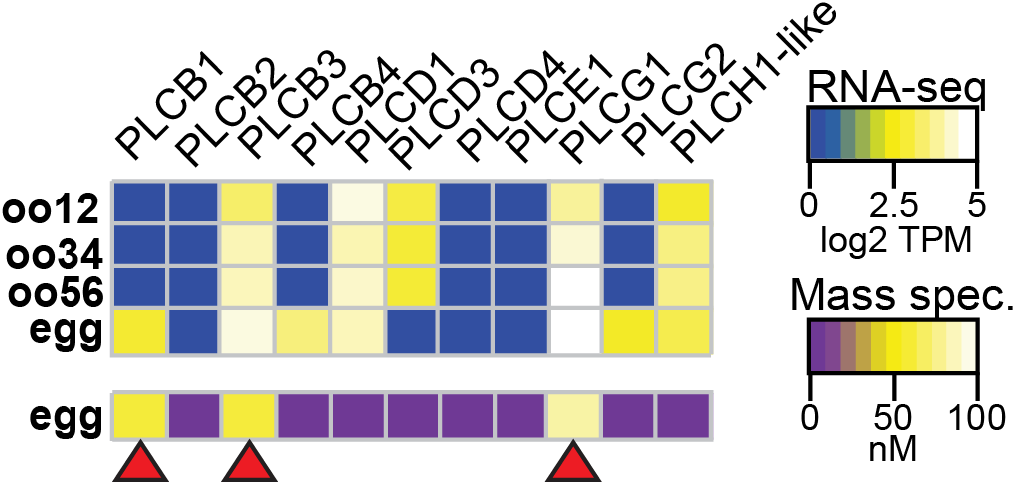
*X. laevis* eggs contain three PLC subtypes. Heatmaps of RNA *(top)* and protein *(bottom)* expression levels of PLC subtypes at varying stages of oocyte and egg development. RNA-seq data displayed as transcripts per million (TPM), and protein data shown as nanomolar concentration. Transcript levels were obtained and compiled from Sessions (2016), while protein concentrations were from Wühr (2014) through mass spectrometry. Red arrows indicate PLC subtypes that are present in both RNA sequencing and mass spectrometry datasets in *X. laevis* eggs.

### Tyrosine phosphorylation of PLCγ1 does not signal the fast block

Typically, PLCγ1 is activated by tyrosine phosphorylation of the critical residue Y776 (homologous to Y783 in mouse PLCγ1) in the SH2 domain of the enzyme (Kadamur and Ross, 2013) (Fig. 3A). Currently, no PLCγ1-specific inhibitors exist, but we can prevent phosphotyrosine-dependent activation of this enzyme using tyrosine kinase inhibitors. Before applying these inhibitors to fertilization, we first screened tyrosine kinase inhibitors for their efficacy in preventing PLCγ1 phosphorylation in *X. laevis* oocytes exogenously expressing the receptor for the platelet derived growth factor (PDGF-R). Applying PDGF to PDGF-R expressing oocytes induced phosphorylation of PLCγ1 at the critical tyrosine at position Y776 (Fig. 3B). We then applied the tyrosine kinase inhibitors, two of which, genistein (100 μM) and dasantinib (100 nM), effectively stopped PDGF-signaled PLCγ1 phosphorylation. By contrast, lavendustin B (100 nM) was only moderately effective, although more than lavendustin A (100 nM) (Fig. 3B).

To determine whether tyrosine phosphorylation of PLCγ1 is required for the fast block, we made whole cell recordings from *X. laevis* eggs inseminated in the presence of the validated tyrosine kinase inhibitors genistein and dasantinib. We did not observe any significant differences between fertilization-evoked depolarizations recorded under control conditions (Fig. 1) or in solutions supplemented with 100 μM genistein or 100 nM dasantinib (Fig. 3C & E). Like eggs fertilized under control conditions, the mean resting potential of eggs in 100 μM genistein was −17.6 ± 1.9 mV and the membrane potential following fertilization was 2.4 ± 2.6 mV (N=7 eggs in 3 independent trials, Fig. 3F & G). The average rate of depolarization was 8.9 ± 3.1 mV/ms (Fig. 3D), similar to the rate recorded under control conditions (4.4 ±1.7 mV, Fig. 1). Eggs recorded in 100 nM dasantinib had a mean resting potential of −10.3 ± 2.2 mV (Fig. 3F) and fertilization potential of −1.3 ± 1.9 mV (Fig. 3G). The rate of depolarization for these eggs was 3.5 ± 1.9 mV/ms (Fig. 3D).

**Figure 3.**
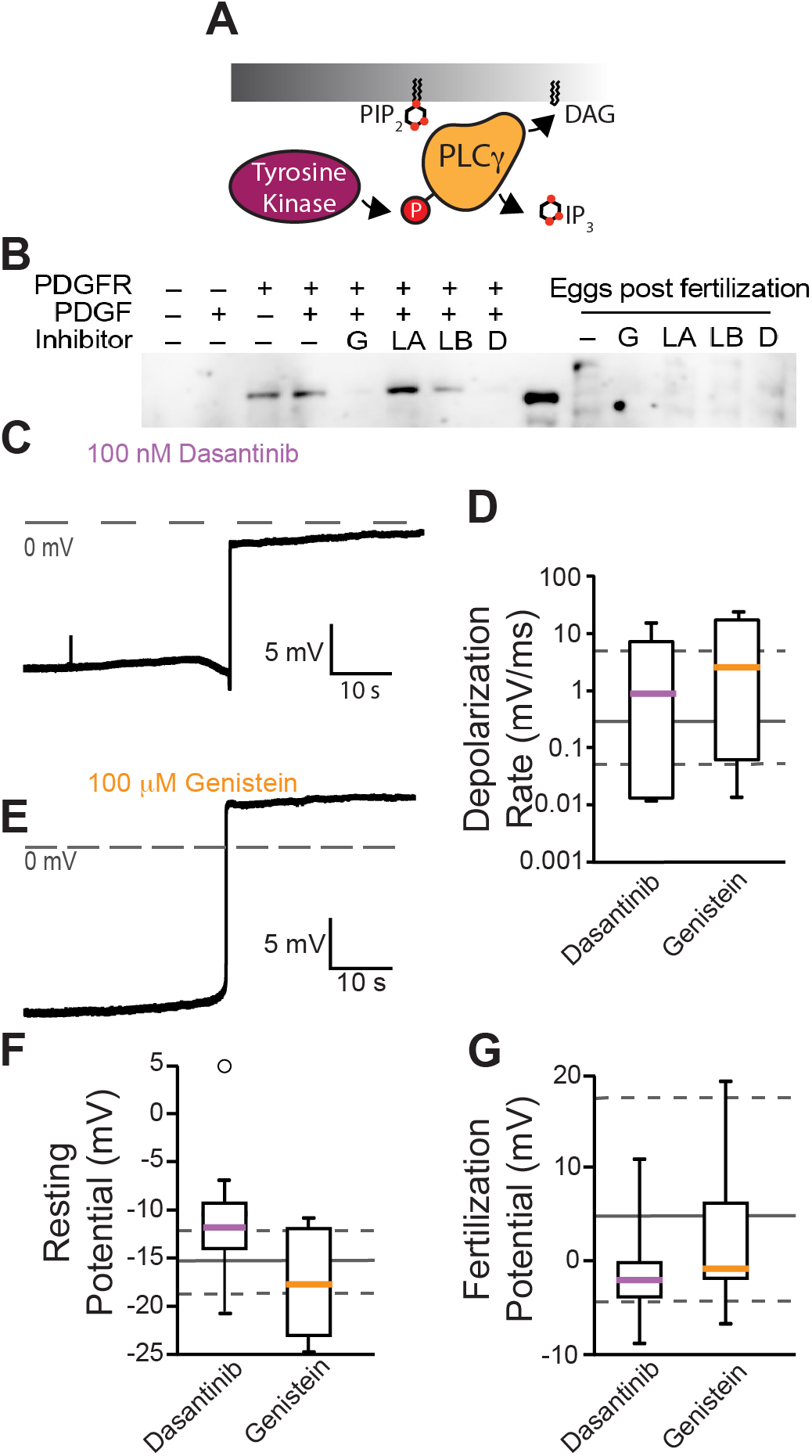
Inhibiting tyrosine phosphorylation of PLCγ1 did not alter the fast block to polyspermy. **(A)** Tyrosine kinases canonically activate PLCγthrough phosphorylation. **(B)** Western blot probing for tyrosine phosphorylation of PLCγ1-Y776 in *X. laevis* oocytes expressing platelet-derived growth factor receptors (PDGFR). Blot probed for PLCγ1-Y776 phosphorylation in the presence of tyrosine kinase inhibitors (G: Genistein, LA: Lavendustin A, LB: Lavendustin B, D: Dasantinib). *X. laevis* eggs were fertilized and processed for western blot revealed that PLCγ1-Y776 was not phosphorylated following fertilization. Representative whole-cell recording of *X. laevis* eggs fertilized in the presence of **(C)** 100 nM dasantinib or **(D)** 100 μM genistein. Tukey box plot distributions of: depolarization rates **(E)**, resting potential **(F)**, and fertilization potential **(G)** in dasantinib and genistein. Middle line denotes the median value, the bIx indicates 25-75%, and the whiskers indicate 10-90%. Gray solid lines indicate median control values while gray dashed lines indicate the 25-75% spread of the controls.

Although lavendustin A and B had little effect on phosphorylation of PLCγ1, others have reported that these compounds stopped the fast block in *X. laevis* fertilization (Glahn et al., 1999). Thus, we made whole cell recordings from *X. laevis* eggs in the presence of 100 nM lavendustin A or lavendustin B. We observed similar resting potentials, −7.9 ± 2.4 mV and −8.1 ± 2.5 mV, and fertilization potentials, 19.7 ± 3.9 mV and 19.8 ± 3.8 mV, in the presence of lavendustin A or B respectively (Fig. S1). We also recorded normal depolarizations in either compound, 0.7 ± 0.5 mV/ms and 0.3 ± 0.2 mV/ms, which were similar to the lavendustin A and B control rates of 0.3 ± 0.2 mV/ms and 0.03 ± 0.01 mV/ms, respectively. Persistence of normal fertilization-evoked depolarizations of eggs inseminated in lavendustin A or B did not disrupt the fast block to polyspermy in *X. laevis* eggs.

To determine whether tyrosine phosphorylation of PLCγ1 occurs during *X. laevis* fertilization, we also blotted for Y776 phosphorylation in *X. laevis* zygotes. *X. laevis* eggs were incubated for 5 min in either control MR/3 conditions or tyrosine kinase inhibitors prior to sperm addition. 10 mins following insemination, we observed contraction of the animal pole, an indicator of fertilization, and eggs were then de-jellied. Following jelly removal, eggs were lysed and processed for western blots as described previously. In 3 independent trials, we observed no evidence of PLCγ1 phosphorylation at Y776 at fertilization (Fig. 3B). Our results demonstrate that PLCγ1 is not activated by tyrosine phosphorylation during *X. laevis* fertilization.

### The *X. laevis* fast block does not require Gα_11_ activation of PLCβ

We next examined the other PLC isoforms present in the egg and investigated whether activation of PLCβis required to signal the fast block. Two types of PLCβs are found in *X. laevis*eggs, PLCβ1 and PLCβ3 (Fig. 2), and these isoforms are typically activated by the α subunit of the Gα_q_ family (Smrcka et al., 1991; Rhee, 2001) (Fig. 4A). The Gα_q_ family is comprised of four members: Gα_q_, Gα_11_, Gα_14_, and Gα_15_ (Peavy et al., 2005). To determine which Gα_q_ family members are expressed in *X. laevis* eggs, we again looked at a proteomics dataset for *X. laevis* eggs (Wuhr et al., 2014). Of the four Gα_q_ subtypes (genes: GNA11, GNA14, GNA15, and GNAQ), we found that only GNA11 is present in the unfertilized *X. laevis* egg (Fig. 4B). We also validated that indeed RNA encoding the Gα_11_ protein was present in the developing oocyte (Fig. 4B).

**Figure 4.**
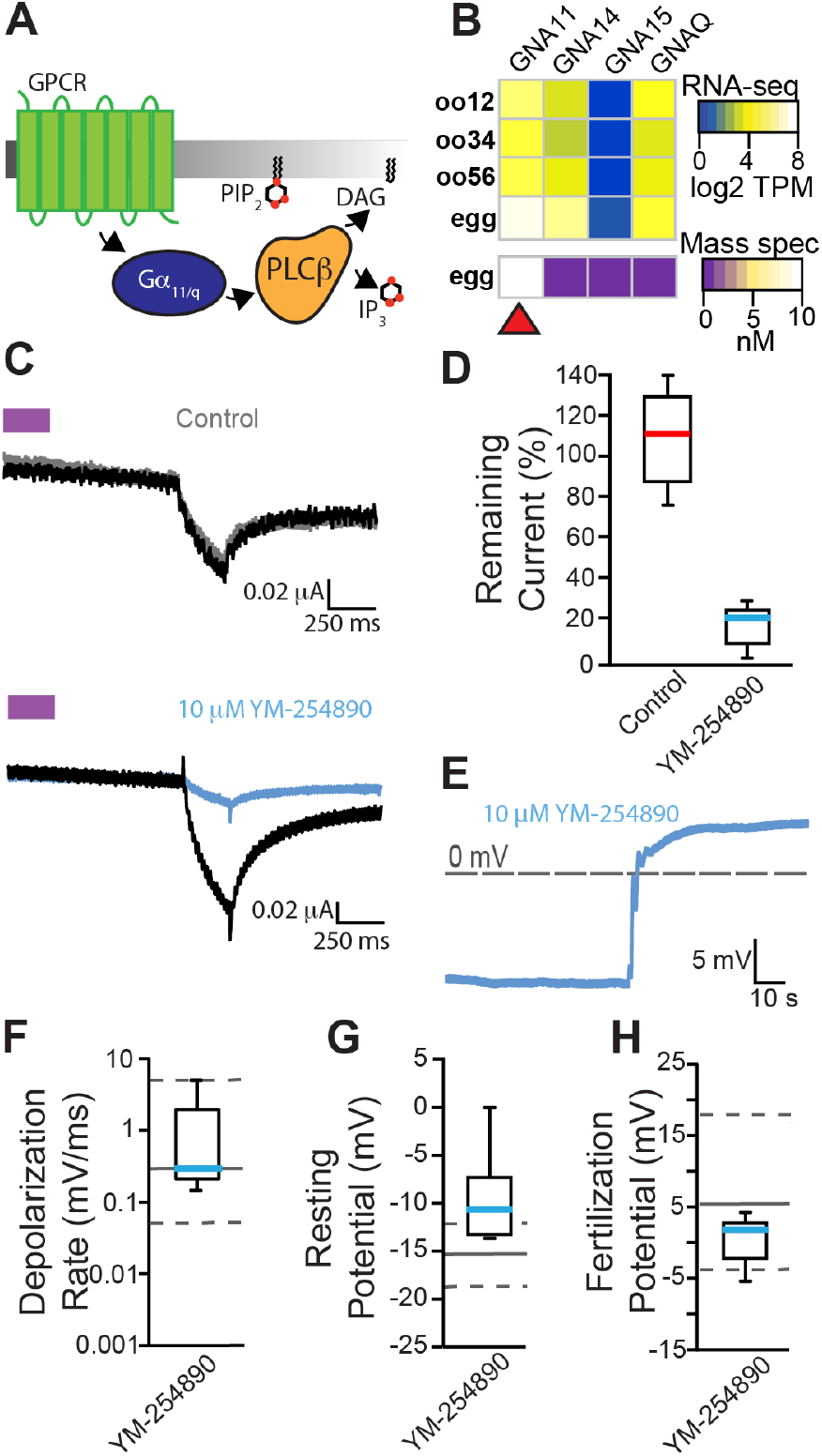
Inhibiting Gα_11/q_ activation of PLCβ did not alter the fast block to polyspermy. **(A)** PLCβ is canonically activated through Gα_11/q_ subunit of G-protein coupled receptors. **(B)** Heatmap of RNA (top) and protein (bottom) expression for the four Gα_q_ family isoforms in *X. laevis* oocyte and fertilization competent eggs. Red arrow indicates the only Gα subtype that was present in both RNA sequencing and mass spectrometry datasets in *X. laevis* eggs: Gα_11_ (encoded by GNA11). **(C)** Representative consecutive two-electrode voltage clamp recordings in control (*top*) conditions in oocytes expressing opto-M1R and clamped at −80 mV, and before and after a 10-minute incubation in 10 μM YM-254890, a Gα_11α11/q_ inhibitor *(bottom)*. The purple block indicates the time of blue/green light application. **(D)** Tukey box plot distributions of the percent remaining current in consecutive recordings in control or 10 μM YM-254890 conditions from oocytes expressing opto-M1R. Middle line indicates median value while the box denotes 25-75% and the whiskers maximum and minimum values. **(E)** Representative whole-cell recording of *X. laevis* eggs in the presence of 10 μM YM-254890. Tukey box plot distributions of **(F)** depolarization rates, **(G)** resting potential, and **(H)** fertilization potential in the presence of YM-254890. Middle line indicates median value while the box denotes 25-75% and the whiskers 10-90%. Gray solid lines denote control median, and dashed gray lines indicate 25-75% distribution of the control.

To identify inhibitors that stop Gα_11_ activation of PLCβ, we made two-electrode voltage clamp recordings from *X. laevis* oocytes expressing the chimeric light-activated muscarinic opto-M1R (Morri et al., 2018). Blue/green light turns on opto-M1R, which then activates PLCβ via Gα_q/11_ and induces IP_3_-mediated Ca^2+^ release from the ER. This burst of intracellular Ca^2+^ is sufficient to activate TMEM16A channels at the membrane, and we can monitor this pathway by observing TMEM16A-conducted currents on *X. laevis* oocytes clamped at −80 mV. We first established that under control conditions, multiple light applications evoked similar amplitudes of TMEM16A conducted currents (Fig. 4C, top). We then screened inhibitors of this pathway by comparing TMEM16A conducted currents before or during application of the known Gα_q/11_ inhibitors YM-254890 (Fig. 4C, bottom) (Uemura et al., 2006) and BIM-46187 (Schmitz et al., 2014). Following an initial light application, opto-M1R expressing oocytes were incubated for 10 mins in the presence of either inhibitor before a second application of light. We found that YM-254890 significantly reduced activation of TMEM16A channels by opto-M1R, reducing the remaining current following the second light application from an average of 108 ±10% remaining current under control conditions to 16 ±5% remaining current in 10 μM YM-254890 (Fig. 4D). By contrast, BIM-46187 did not alter opto-M1R activation of TMEM16A conducted current, with 103 ±17% remaining current in 25 μM BIM-46187 (Fig. S2). These data indicate that YM-254890 effectively stops Gα_q/11_ activation of PLCβ.

To test whether Gα_11_ activates PLCβ during the fast block, we performed whole cell recordings on *X. laevis* eggs in the presence of 10 μM YM-254890 following a ten-minute incubation. In six independent trials, we observed normal fertilization-evoked depolarizations in the presence of 10 μM YM-254890 (Fig. 4E). The average membrane potentials from eggs inseminated in 10 μM YM-254890 was −9.3 ± 1.9 mV before and 0.5 ± 1.3 mV after fertilization (Fig. 4G & H). The depolarization rates were also similar in the presence of absence of YM-254890, with an average 1.1 ± 0.7 mV/ms (Fig. 4F). These findings reveal that fertilization does not require activation of a Gα_11_ coupled activation of PLCβ.

## Discussion

The fast block to polyspermy is one of the earliest processes used by external fertilizers to ensure the genetic integrity of the nascent zygote. Although this process is used by the diverse assortment of animals that fertilize outside of the mother, the signaling events involved in this process have largely remained elusive. Our data demonstrate that the fast block is mediated by activation of a PLC (Fig. 1). Specifically, we have found that in the presence of the general PLC inhibitor U73122, fertilization did not depolarize *X. laevis* eggs, and that these eggs developed asymmetric cleavage furrows (Fig. 1F), an indicator of polyspermy. We have previously reported that the fast block in *X. laevis* eggs requires an activation of the ER-localized IP3R (Wozniak et al., 2018b), which enables a Ca^2+^ release and opening of the Ca^2+^ activated Cl^−^ channel TMEM16A (Wozniak et al., 2018a). Cl^−^ then leaves the egg to depolarize the plasma membrane (Cross and Elinson, 1980; Grey et al., 1982; Webb and Nuccitelli, 1985). Here we sought to uncover how fertilization initiates the fast block in *X. laevis* eggs by examining how fertilization activates PLC.

We used proteomic (Wuhr et al., 2014) and transcriptomic (Session et al., 2016) data on PLC isoforms present in the *X. laevis* eggs to provide clues about how they are activated by fertilization. Interrogating these datasets, we identified three PLC isoforms in *X. laevis* eggs: PLCγ1, PLCβ1 and PLCβ3. Notably, PLCγ1 is approximately 20-fold more abundant than either PLCβ1 or PLCβ3 (Fig. 2). PLC subfamilies are distinguished by structural features that serve as regulatory elements for cell signaling pathways. However, the catalytic component of PLC enzymes is highly conserved between members of the distinct PLC subfamilies (Gresset et al., 2012). Consequently, there are no inhibitors specific to each PLC subtype. However, selective inhibition can be accomplished upstream of PLC activation *e.g*. the kinase and GPCR-coupled mechanisms that activate PLCγ and PLCβ, respectively).

The PLCγ enzyme is typically activated by phosphorylation of a critical tyrosine residue near its catalytic domain (Gresset et al., 2012). We validated that the tyrosine kinase inhibitors genistein and dasantanib blocked PLCγ1 phosphorylation by the receptor tyrosine kinase PDGF-R, but found that neither had any effect on the fast block (Fig. 3B & D). These data indicate that tyrosine kinase-induced activation of PLCγ1 does not mediate the fast block in *X. laevis*.

Intriguingly, our data disagree with the prevailing hypothesis that fertilization in *X. laevis* activates PLCγ1 via tyrosine phosphorylation (Mahbub Hasan et al., 2005). In this hypothesis, a Src family kinase tyrosine phosphorylates PLCγ to thereby activate increased intracellular Ca^2+^ that initiates the cortical granule exocytosis of the slow block to polyspermy (Sato et al., 2000). Although it is yet to be determined whether the same signaling pathway initiates both the fast and slow block, our data suggests that fertilization does not activate PLCγ1 via tyrosine phosphorylation of Y776 (Fig. 3B). We suspect that differences in the methods used to monitor tyrosine phosphorylation of PLCγ1 following fertilization underlies the differences in these data.

We also considered a possible role for a G-protein mediated signaling pathway in the *X. laevis* fast block. Typically, PLCβ isoforms are activated by G-protein pathways requiring members of the Gα_q/11_ family. We noted that *X. laevis* eggs express the Gα_11_ subunit, supporting this possibility. By activating Gα_q/11_ through the light-activated receptor opto-M1R (Morri et al., 2018), we were able to reproducibly induce TMEM16A currents and furthermore inhibit this effect by application of the Gα_q/11_ inhibitor YM-254890. However, application of 10 μM YM-254890 had no effect on the fast block to polyspermy in *X. laevis* eggs (Fig. 4B).

In addition to Gα_q/11_, PLCβ subtypes can be activated by the Gβγ dimers derived from Gα_i_ signaling pathways (Boyer et al., 1992). Generally, Gα_i_ is the most abundant a subunit isoform that is ubiquitously expressed in all cells, and consequently the most prolific source of Gβγ signaling (Hepler and Gilman, 1992). Even though all G-protein heterotrimers contain β and γ subunits, initiation of Gβγ signaling downstream of G-protein coupled receptors is typically the result of Gα_i_ coupled receptors, likely due to the relatively low affinity of Gβγ for its effectors relative to interactions of Gα subunits and increased cellular abundance of Gα_i_ heterotrimers (Hepler and Gilman, 1992). Indeed, *X. laevis* eggs express three Gα_i_ isoforms abundantly (Fig. S3). However we did not pursue a role for Gα_i_ because published data suggest that this pathway did not contribute to the fast block (Kline et al., 1991). Specifically, Kline and colleagues demonstrated that stopping Gα_i_ signaling with pertussis toxin, did not alter the fertilization signaled depolarization in *X. laevis* eggs (Kline et al., 1991).

Our data agrees with interpretations of results published by Runft and colleagues, who loaded *X. laevis* eggs with calcium green dextran to observe the fertilization signaled Ca^2+^ wave (Runft et al., 1999). They reported that stopping PLCγ1 activation by expressing a dominant negative SH2 domain did not alter the fertilization activated Ca^2+^ wave in *X. laevis* eggs (Runft et al., 1999). The SH2 domain includes the Y783 residue within the enzyme’s active site that mediates activation via phosphorylation (Hajicek et al., 2019). We did not observe evidence that this residue is phosphorylated, leaving the possibility that overexpression of the SH2 domain does not interfere with a phosphorylation-independent mechanism for PLCγ1 activation. Runft and colleagues also reported that injection of a Gα_q_ targeting antibody did not disrupt the fertilization signaled Ca^2+^ wave. *X. laevis* eggs don’t express the Gα_q_ protein, but antibody injection was sufficient to disrupt PLCβ-mediated Ca^2+^ via an exogenously expressed serotonin receptor, suggesting that their antibody was capable of inhibiting the native Gα_q/11_ protein (Runft et al., 1999).

Our data supports the hypothesis where a non-canonical pathway activates a PLC to signal the fast block in *X. laevis* eggs. A possible mechanism for this is that elevated cytoplasmic Ca^2+^ alone may signal PLC activation at fertilization. Elevated Ca^2+^ has been shown to activated several PLC isoforms, including PLCγ1 (Wahl et al., 1992; Hwang et al., 1996) and PLCβ (Ryu et al., 1987; Wahl et al., 1992; Hwang et al., 1996). Several groups have proposed that this mechanism underlies the regenerative Ca^2+^ wave that enables cortical granule exocytosis for the slow polyspermy block (Fall et al., 2004; Wagner et al., 2004).

Alternatively, a sperm donated PLC could give rise to the fast block. In mammalian fertilization, the sperm-derived, soluble PLCζ is released as a consequence of sperm entry. Released PLCζ signals the Ca^2+^ wave and initiates the slow polyspermy block (Nozawa et al., 2018). However, the PLCZ gene has not yet been annotated in non-mammalian animals. Whether a different sperm derived PLC activates eggs is yet to be determined.

Fertilization quickly activates a PLC to signal TMEM16A activation and the fast block to polyspermy in *X. laevis*. Several questions remain regarding this critical process. Answering these questions will not only reveal the earliest events of new life but will also shed light on the voltage-dependence of fertilization.

## Acknowledgments

We thank D. Summerville and P. Sau for excellent technical assistance. This work was supported by National Institute for Health grant 1R01GM125638 to A.E.C.

## Cited References

Bianchi, E., and G.J. Wright. 2016. Sperm Meets Egg: The Genetics of Mammalian Fertilization. Annu Rev Genet. 50:93–111.

Boyer, J.L., G.L. Waldo, and T.K. Harden. 1992. Beta gamma-subunit activation of G-protein-regulated phospholipase C. J Biol Chem. 267:25451–25456.

Cross, N.L., and R.P. Elinson. 1980. A fast block to polyspermy in frogs mediated by changes in the membrane potential. Dev Biol. 75:187–198.

Elinson, R.P. 1975. Site of sperm entry and a cortical contraction associated with egg activation in the frog Rana pipiens. Dev Biol. 47:257–268.

Evans, J.P. 2020. Preventing polyspermy in mammalian eggs-Contributions of the membrane block and other mechanisms. Mol Reprod Dev. 87:341–349.

Fahrenkamp, E., B. Algarra, and L. Jovine. 2020. Mammalian egg coat modifications and the block to polyspermy. Mol Reprod Dev. 87:326–340.

Fall, C.P., J.M. Wagner, L.M. Loew, and R. Nuccitelli. 2004. Cortically restricted production of IP3 leads to propagation of the fertilization Ca2+ wave along the cell surface in a model of the Xenopus egg. J Theor Biol. 231:487–496.

Fontanilla, R.A., and R. Nuccitelli. 1998. Characterization of the sperm-induced calcium wave in *Xenopus* eggs using confocal microscopy. Biophys J. 75:2079–2087.

Gagoski, D., M.E. Polinkovsky, S. Mureev, A. Kunert, W. Johnston, Y. Gambin, and K. Alexandrov. 2016. Performance benchmarking of four cell-free protein expression systems. Biotechnol Bioeng. 113:292–300.

Glahn, D., S.D. Mark, R.K. Behr, and R. Nuccitelli. 1999. Tyrosine kinase inhibitors block sperm-induced egg activation in Xenopus laevis. Dev Biol. 205:171–180.

Gresset, A., J. Sondek, and T.K. Harden. 2012. The phospholipase C isozymes and their regulation. Subcell Biochem. 58:61–94.

Grey, R.D., M.J. Bastiani, D.J. Webb, and E.R. Schertel. 1982. An electrical block is required to prevent polyspermy in eggs fertilized by natural mating of Xenopus laevis. Dev Biol. 89:475–484.

Hajicek, N., N.C. Keith, E. Siraliev-Perez, B.R. Temple, W. Huang, Q. Zhang, T.K. Harden, and J. Sondek. 2019. Structural basis for the activation of PLC-gamma isozymes by phosphorylation and cancer-associated mutations. Elife. 8.

Hassold, T., N. Chen, J. Funkhouser, T. Jooss, B. Manuel, J. Matsuura, A. Matsuyama, C. Wilson, J.A. Yamane, and P.A. Jacobs. 1980. A cytogenetic study of 1000 spontaneous abortions. Ann Hum Genet. 44:151–178.

Hepler, J.R., and A.G. Gilman. 1992. G proteins. Trends in biochemical sciences. 17:383–387.

Hill, W.G., N.M. Southern, B. MacIver, E. Potter, G. Apodaca, C.P. Smith, and M.L. Zeidel. 2005. Isolation and characterization of the Xenopus oocyte plasma membrane: a new method for studying activity of water and solute transporters. Am J Physiol Renal Physiol. 289:F217–224.

Hwang, S.C., D.Y. Jhon, Y.S. Bae, J.H. Kim, and S.G. Rhee. 1996. Activation of phospholipase C-gamma by the concerted action of tau proteins and arachidonic acid. J Biol Chem. 271:18342–18349.

Jaffe, L.A. 1976. Fast block to polyspermy in sea urchin eggs is electrically mediated. Nature. 261:68–71.

Jaffe, L.A., and M. Gould. 1985. Polyspermy-preventing mechanisms. In Biology of Fertilization. C.B. Metz and A. Monroy, editors. Academic Press, New York. 223–250.

Kadamur, G., and E.M. Ross. 2013. Mammalian phospholipase C. Annu Rev Physiol. 75:127–154.

Kline, D., G.S. Kopf, L.F. Muncy, and L.A. Jaffe. 1991. Evidence for the involvement of a pertussis toxin-insensitive G-protein in egg activation of the frog, Xenopus laevis. Dev Biol. 143:218–229.

Mahbub Hasan, A.K., K. Sato, K. Sakakibara, Z. Ou, T. Iwasaki, Y. Ueda, and Y. Fukami. 2005. Uroplakin III, a novel Src substrate in Xenopus egg rafts, is a target for sperm protease essential for fertilization. Dev Biol. 286:483–492.

Morri, M., I. Sanchez-Romero, A.M. Tichy, S. Kainrath, E.J. Gerrard, P.P. Hirschfeld, J. Schwarz, and H. Janovjak. 2018. Optical functionalization of human Class A orphan G-protein-coupled receptors. Nat Commun. 9:1950.

Nozawa, K., Y. Satouh, T. Fujimoto, A. Oji, and M. Ikawa. 2018. Sperm-borne phospholipase C zeta-1 ensures monospermic fertilization in mice. Sci Rep. 8:1315.

Nuccitelli, R., D.L. Yim, and T. Smart. 1993. The sperm-induced Ca2+ wave following fertilization of the Xenopus egg requires the production of Ins(1, 4, 5)P3. Dev Biol. 158:200–212.

Peavy, R.D., K.B. Hubbard, A. Lau, R.B. Fields, K. Xu, C.J. Lee, T.T. Lee, K. Gernert, T.J. Murphy, and J.R. Hepler. 2005. Differential effects of Gq alpha, G14 alpha, and G15 alpha on vascular smooth muscle cell survival and gene expression profiles. Mol Pharmacol. 67:2102–2114.

Rhee, S.G. 2001. Regulation of phosphoinositide-specific phospholipase C. Annu Rev Biochem. 70:281–312.

Rojas, J., F. Hinostroza, S. Vergara, I. Pinto-Borguero, F. Aguilera, R. Fuentes, and I. Carvacho. 2021. Knockin’ on Egg’s Door: Maternal Control of Egg Activation That Influences Cortical Granule Exocytosis in Animal Species. Front Cell Dev Biol. 9:704867.

Runft, L.L., J. Watras, and L.A. Jaffe. 1999. Calcium release at fertilization of *Xenopus* eggs requires type I IP_3_ receptors, but not SH2 domain-mediated activation of PLCγ or G_q_- mediated activation of PLCβ. Dev Biol. 214:399–411.

Ryu, S.H., P.G. Suh, K.S. Cho, K.Y. Lee, and S.G. Rhee. 1987. Bovine brain cytosol contains three immunologically distinct forms of inositolphospholipid-specific phospholipase C. Proc Natl Acad Sci U S A. 84:6649–6653.

Sato, K., A.A. Tokmakov, T. Iwasaki, and Y. Fukami. 2000. Tyrosine kinase-dependent activation of phospholipase Cgamma is required for calcium transient in Xenopus egg fertilization. Dev Biol. 224:453–469.

Schmitz, A.L., R. Schrage, E. Gaffal, T.H. Charpentier, J. Wiest, G. Hiltensperger, J. Morschel, S. Hennen, D. Haussler, V. Horn, D. Wenzel, M. Grundmann, K.M. Bullesbach, R. Schroder, H.H. Brewitz, J. Schmidt, J. Gomeza, C. Gales, B.K. Fleischmann, T. Tuting, D. Imhof, D. Tietze, M. Gutschow, U. Holzgrabe, J. Sondek, T.K. Harden, K. Mohr, and E. Kostenis. 2014. A cell-permeable inhibitor to trap Galphaq proteins in the empty pocket conformation. Chem Biol. 21:890–902.

Session, A.M., Y. Uno, T. Kwon, J.A. Chapman, A. Toyoda, S. Takahashi, A. Fukui, A. Hikosaka, A. Suzuki, M. Kondo, S.J. van Heeringen, I. Quigley, S. Heinz, H. Ogino, H. Ochi, U. Hellsten, J.B. Lyons, O. Simakov, N. Putnam, J. Stites, Y. Kuroki, T. Tanaka, T. Michiue, M. Watanabe, O. Bogdanovic, R. Lister, G. Georgiou, S.S. Paranjpe, I. van Kruijsbergen, S. Shu, J. Carlson, T. Kinoshita, Y. Ohta, S. Mawaribuchi, J. Jenkins, J. Grimwood, J. Schmutz, T. Mitros, S.V. Mozaffari, Y. Suzuki, Y. Haramoto, T.S. Yamamoto, C. Takagi, R. Heald, K. Miller, C. Haudenschild, J. Kitzman, T. Nakayama, Y. Izutsu, J. Robert, J. Fortriede, K. Burns, V. Lotay, K. Karimi, Y. Yasuoka, D.S. Dichmann, M.F. Flajnik, D.W. Houston, J. Shendure, L. DuPasquier, P.D. Vize, A.M. Zorn, M. Ito, E.M. Marcotte, J.B. Wallingford, Y. Ito, M. Asashima, N. Ueno, Y. Matsuda, G.J. Veenstra, A. Fujiyama, R.M. Harland, M. Taira, and D.S. Rokhsar. 2016. Genome evolution in the allotetraploid frog Xenopus laevis. Nature. 538:336–343.

Smrcka, A.V., J.R. Hepler, K.O. Brown, and P.C. Sternweis. 1991. Regulation of polyphosphoinositide-specific phospholipase C activity by purified Gq. Science. 251:804–807.

Tembo, M., M.L. Sauer, B.W. Wisner, D.O. Beleny, and M.A. Napolitano. 2020. Inhibiting actin polymerization does not prevent the fast block to polyspermy in the African clawed frog, Xenopus laevis. bioRxiv.

Tembo, M., K.L. Wozniak, R.E. Bainbridge, and A.E. Carlson. 2019. Phosphatidylinositol 4,5- bisphosphate (PIP2) and Ca(2+) are both required to open the Cl(-) channel TMEM16A. J Biol Chem. 294:12556–12564.

Uemura, T., T. Kawasaki, M. Taniguchi, Y. Moritani, K. Hayashi, T. Saito, J. Takasaki, W. Uchida, and K. Miyata. 2006. Biological properties of a specific Galpha q/11 inhibitor, YM-254890, on platelet functions and thrombus formation under high-shear stress. Br J Pharmacol. 148:61–69.

Wagner, J., C.P. Fall, F. Hong, C.E. Sims, N.L. Allbritton, R.A. Fontanilla, Moraru, II, L.M. Loew, and R. Nuccitelli. 2004. A wave of IP3 production accompanies the fertilization Ca2+ wave in the egg of the frog, Xenopus laevis: theoretical and experimental support. Cell Calcium. 35:433–447.

Wahl, M.I., G.A. Jones, S. Nishibe, S.G. Rhee, and G. Carpenter. 1992. Growth factor stimulation of phospholipase C-gamma 1 activity. Comparative properties of control and activated enzymes. J Biol Chem. 267:10447–10456.

Webb, D.J., and R. Nuccitelli. 1985. Fertilization potential and electrical properties of the Xenopus laevis egg. Dev Biol. 107:395–406.

Wong, J.L., and G.M. Wessel. 2006. Defending the zygote: search for the ancestral animal block to polyspermy. Curr Top Dev Biol. 72:1–151.

Wozniak, K.L., R.E. Bainbridge, D.W. Summerville, M. Tembo, W.A. Phelps, M.L. Sauer, B.W. Wisner, M.E. Czekalski, S. Pasumarthy, M.L. Hanson, M.B. Linderman, C.H. Luu, M.E. Boehm, S.M. Sanders, K.M. Buckley, D.J. Bain, M.L. Nicotra, M.T. Lee, and A.E. Carlson. 2020. Zinc protection of fertilized eggs is an ancient feature of sexual reproduction in animals. PLoS Biol. 18:e3000811.

Wozniak, K.L., and A.E. Carlson. 2020. Ion channels and signaling pathways used in the fast polyspermy block. Mol Reprod Dev. 87:350–357.

Wozniak, K.L., B.L. Mayfield, A.M. Duray, M. Tembo, D.O. Beleny, M.A. Napolitano, M.L. Sauer, B.W. Wisner, and A.E. Carlson. 2017. Extracellular Ca2+ Is Required for Fertilization in the African Clawed Frog, Xenopus laevis. PLoS One. 12:e0170405.

Wozniak, K.L., W.A. Phelps, M. Tembo, M.T. Lee, and A.E. Carlson. 2018a. The TMEM16A channel mediates the fast polyspermy block in Xenopus laevis. J Gen Physiol.

Wozniak, K.L., M. Tembo, W.A. Phelps, M.T. Lee, and A.E. Carlson. 2018b. PLC and IP3- evoked Ca(2+) release initiate the fast block to polyspermy in Xenopus laevis eggs. J Gen Physiol.

Wuhr, M., R.M. Freeman, Jr., M. Presler, M.E. Horb, L. Peshkin, S.P. Gygi, and M.W. Kirschner. 2014. Deep proteomics of the Xenopus laevis egg using an mRNA-derived reference database. Curr Biol. 24:1467–1475.

Yang, J., T. Aguero, and M.L. King. 2015. The Xenopus Maternal-to-Zygotic Transition from the Perspective of the Germline. Curr Top Dev Biol. 113:271–303.

