## Supplemental Figure 1 for "TMEM16A activation for the fast block to polyspermy in the African clawed frog does not require conventional activation of egg PLCs"

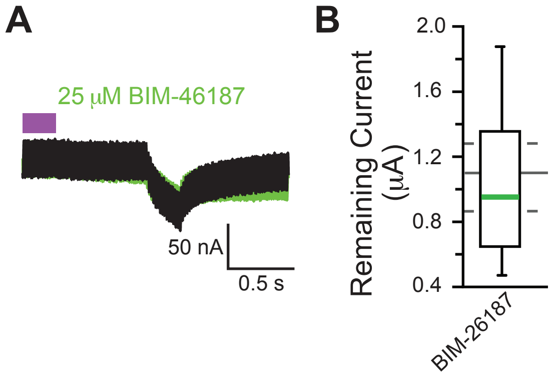


Supplemental Figure 2: **BIM-46187 did not inhibit G$\alpha_{11}$ activation of PLC$\beta$ in *X. laevis* oocytes. (A)** Representative consecutive two-electrode voltage clamp recordings in oocytes expressing opto-M1R and clamped at -80 mV, before and after a 10-minute incubation in 25 μM BIM-46187. The purple block indicates the time of blue/green light application. **(B)** Tukey box plot distributions of the percent remaining current in consecutive recordings in 25 μM BIM-46187 from oocytes expressing opto-M1R. Middle line indicates median value while the box denotes 25-75% and the whiskers maximum and minimum values. The gray solid line represents the median, and the gray dashed lines denote 25-75% distribution, of the data collected under control conditions.
