## Supplemental Figure 3 for "TMEM16A activation for the fast block to polyspermy in the African clawed frog does not require conventional activation of egg PLCs"

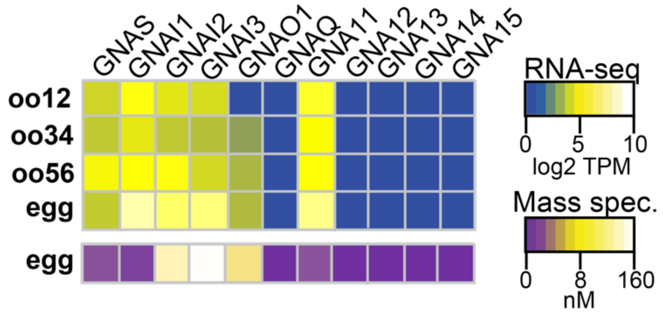


Supplemental Figure 3: **GNA isoforms expressed in *X. laevis* eggs.** Heatmaps of RNA (top) and protein (bottom) expression levels of G$\alpha$ subunits at various developmental stages of egg maturation. RNA-seq data displayed as transcripts per million (TPM), and protein data shown as nanomolar concentration. Transcript levels were obtained and compiled from Sessions (2016), while protein concentrations were from Wühr (2014) through mass spectrometry.
